# DeepDDS: deep graph neural network with attention mechanism to predict synergistic drug combinations

**DOI:** 10.1101/2021.04.06.438723

**Authors:** Jinxian Wang, Xuejun Liu, Siyuan Shen, Lei Deng, Hui Liu

## Abstract

**Motivation:** Drug combination therapy has become a increasingly promising method in the treatment of cancer. However, the number of possible drug combinations is so huge that it is hard to screen synergistic drug combinations through wet-lab experiments. Therefore, computational screening has become an important way to prioritize drug combinations. Graph neural network have recently shown remarkable performance in the prediction of compound-protein interactions, but it has not been applied to the screening of drug combinations.

**Results:** In this paper, we proposed a deep learning model based on graph neural networks and attention mechanism to identify drug combinations that can effectively inhibit the viability of specific cancer cells. The feature embeddings of drug molecule structure and gene expression profiles were taken as input to multi-layer feedforward neural network to identify the synergistic drug combinations. We compared DeepDDS with classical machine learning methods and other deep learning-based methods on benchmark data set, and the leave-one-out experimental results showed that DeepDDS achieved better performance than competitive methods. Also, on an independent test set released by well-known pharmaceutical enterprise AstraZeneca, DeepDDS was superior to competitive methods by more than 16% predictive precision. Furthermore, we explored the interpretability of the graph attention network, and found the correlation matrix of atomic features revealed important chemical substructures of drugs. We believed that DeepDDS is an effective tool that prioritized synergistic drug combinations for further wet-lab experiment validation.

**Availability and implementation:** Source code and data are available at https://github.com/Sinwang404/DeepDDS/tree/master

## Introduction

Both traditional and modern medicine has taken advantage of the combined use of several active agents to treat diseases. Compared with single-drug therapy, the drug combinations often improve efficacy (Csermely et al., 2013), reduce side effects (Zhao et al., 2013) and overcome drug resistance (Hill et al., 2013, Verderosa et al.). Drug combinations are increasingly used to treat a variety of complex diseases, such as hypertension (Giles et al., 2014), infectious diseases (Zheng et al., 2018), and cancer (Kim et al., 2021; Vitiello et al., 2021). For example, triple-negative breast cancer is a malignant tumor with strong invasiveness, high metastasis rate and poor prognosis. Lapatinib or Rapamycin alone has little therapeutic effect, but their combined treatment has been reported to significantly increase the apoptosis rate of triple-negative breast cancer cells (Liu et al., 2011). However, some drug combinations may cause antagonistic effect and even aggravate the disease (Azam and Vazquez, 2021). Therefore, it is crucial to accurately discover synergistic drug combinations to specific diseases.

Traditional discovery of drug combinations is mainly based on clinical trials and limited to only a few number of drugs (Li et al., 2015), far from meeting the urgent need for anticancer drugs. Due to the great number of possible drug combinations, traditional method is cost-consuming and impractical. With the development of high-throughput drug screening technology, people can simultaneously carry out large-scale screening of drug combinations over hundreds of cancer cell lines (Hertzberg and Pope, 2000, Bajorath, 2002, Macarron et al., 2011). Torres *et al*. utilized yeast to screen a large number of drug combinations and provided a method to identify preferential drug combinations for further testing in human cells (Torres et al., 2013). In despite of high degree of genomic correlation between the original tumor and the derived cancer cell line, *in vitro* experiments of high-throughput drug screening still cannot accurately capture the mode of action of drug molecules *in vivo* (Ferreira et al., 2013). Microcalorimetry screening (Kragh et al., 2021) and genetically encoded fluorescent sensors (Potekhina et al., 2021) have been developed to screen effective antimicrobial combinations for *in vivo* disease treatment. However, these techniques require skilled operations and complicated experimental procedures.

In recent years, the datasets of single drug sensitivities to cancer cell lines increase greatly, such as Cancer Cell Line Encyclopedia (CCLE) (Barretina et al., 2012) and Genomics of Drug Sensitivity in Cancer (GDSC), which contains drug sensitivities to hundreds of human cancer cell lines, as well as gene expression profiles, mutants and copy number variants. Meanwhile, several large-scale data resource of drug combinations have been released. For example, O’Neil *et al*. released a large-scale drug pair synergy study, which included more than 20,000 pairwise synergy scores between 38 unique drugs (O’Neil et al., 2016). The famous pharmaceutical company AstraZeneca (Menden et al., 2019) released their drug pair collaboration experiments, which includes 11,576 experiments of 910 drug combinations to 85 cancer cell lines with genome-related information. DrugCombDB (Liu et al., 2020) has collected more than 6,000,000 quantitative drug dose responses, by which they calculated synergy scores to evaluate synergy or antagonism for each drug combination. In addition, quite a few data portal designed to collect drug combinations and relevant knowledge have been developed. The release of above-mentioned data resources motivated the development of computational screening of drug combinations. Many studies have been proposed to explore the vast space of drug combinations to identify synergistic efficacy. For example, classical machine learning methods, such as support vector machine (SVM) and random forest, successfully predicted the maximal antiallodynic effect of a new derivative of dihydrofuran-2-one (LPP1) used in combination with pregabalin (PGB) in the streptozocin-induced neuropathic pain model in mice (Sałat and Sałat, 2013, Qi, 2012).

Recently, the deep learning is increasingly applied to drug development and discovery. For example, DeepSynergy (Preuer et al., 2018) combined the chemical information of drugs and genomic features of cancer cells to predict drug pairs with synergistic effects. TranSynergy (Liu and Xie, 2021) is a mechanism-driven and self-attention boosted deep learning model that integrates information from gene-gene interaction networks, gene dependencies, and drug-target associations to predict synergistic drug combinations and deconvolute the cellular mechanisms. On the other hand, some studies applied SMILES to characterize chemical properties of drugs. For example, Gao et al. used the drug descriptors based on the SMILES to predict drug synergy. Liu et al. regarded the SMILES code as a string and directly input into a convolutional neural network (Liu et al., 2016) to extract drug features for subsequent prediction task. More interesting, graph neural network is used to learn feature representation from drug chemical structure (Wu et al., 2018, Xiong et al., 2019).

In this paper, we propose a deep learning model, DeepDDS (Deep Learning for Drug-Drug Synergy prediction), to predict the synergistic effect of drug combinations. First, the drug chemical structure is represented by a graph in which the vertices are atoms and the edges are chemical bonds. Next, a graph convolutional network and attention mechanism is used to compute the drug embedding vectors. By integration of the genomic and pharmaceutical features, DeepDDS can capture important information from drug chemical structure and gene expression patterns to identify synergistic drug combinations to specific cancer cell lines. We compare DeepDDS to both classical machine learning methods (SVM, RF, GTB and XGBoost) and other latest deep learning (DTF, DeepSynergy and TranSynergy) on benchmark data set, DeepDDS significantly outperform other competitive methods. In particular, we conducted leave-one-out experiments to verify that DeepDDS achieved better performance when one drug (combination) or one tissue is not included in the training set. Also, on an independent test set released by well-known pharmaceutical enterprise AstraZeneca, DeepDDS was superior to competitive methods by more than 16% predictive precision. We also explored the function of graph attention network in revealing important chemical substructures of drugs, and found the correlation matrix of atomic features showed clustering patterns among atom subgroups during the training process. Finally, we use the trained model to predict novel drug combination and find two previously reported synergistic drug combinations in the top 10 predicted results, MK2206 and AZD5363, MK2206 and AZD6244 to HCC1806 breast cancer cells. In summary, we believed that DeepDDS is an effective tool that prioritized synergistic drug combinations for further wet-lab experiment validation.

## Materials and methods

### Data source

The SMILES (Simplified Molecular Input Line Entry System) (Weininger, 1988) of drugs are obtained from DrugBank ([Wishart et al., 2018]), based on which the chemical structure of a drug can be converted to a graph using RDKit ([Landrum et al., 2006]). In the molecular graph, the vertices are atoms and the edges are chemical bonds.

The gene expression data of cancer cell lines are obtained from Cancer Cell Line Encyclopedia (CCLE, Barretina et al., 2012), which is an independent project that makes effort to characterize genomes, mRNA expression, and anti-cancer drug dose responses across cancer cell lines. The expression data is normalized through TPM (Transcripts Per Million) based on the genome-wide read counts matrix.

To construct the benchmark set, we obtain the drug combination sensitivity data from a recently released largescale oncology screening data set (O’Neil et al., 2016), where the viability of 39 cancer cells treated with thousands of drug combinations was evaluated by biochemical assay. The Loewe Additivity score (Loewe, 1953), a quantitative metric that defines the synergistic or antagonistic effect of the drug combination, was calculated based on the 4 by 4 doseresponse matrix using the Combenefit tool (Di Veroli et al., 2016). Of note, multiple replicates of one drug combination were assayed in the original data, and thus the average score of the replicates was selected as the final synergistic score for each unique drug-pair-cell-line combination. According to the Loewe score, a combination with the score above zero is regarded as synergistic, and with the score below zero is antagonistic. Obviously, the drug combinations with higher synergistic scores are more attractive candidates for further clinical experiments. Since many additive combinations may exist (synergy scores are around 0 due to noise), we choose a stricter threshold to classify the combinations. Particularly, combinations with synergy score higher than 10 are labeled as positive (synergistic), and those with score less than 0 are labeled as negative (antagonistic). This yielded a balanced benchmark set that contains 12,415 unique drug pair-cell line combinations, covering 36 anticancer drugs and 31 human cancer cell lines.

### Pipeline of DeepDDS

Figure 1 illustrates the end-to-end learning framework for the prediction of synergistic drug combinations. For each pairwise drug combination, the input layer firstly receives the molecular graphs of two drugs and gene expression profiles of one cancer cell line that was treated by these two drugs. We tested two type of Graph Neural Networks (GNN), graph attention network (GAT) and graph convolution network (GCN), to extract features of drugs. The genomic feature representation of cancer cells is encoded by a multi-layer perception (MLP). The embedding vectors are subsequently concatenated as the final feature representation of each drug-pair-cell-line combination, which is propagated through the fully-connected layers for the binary classification of drug combinations (synergistic or antagonistic).

**Fig. 1.**
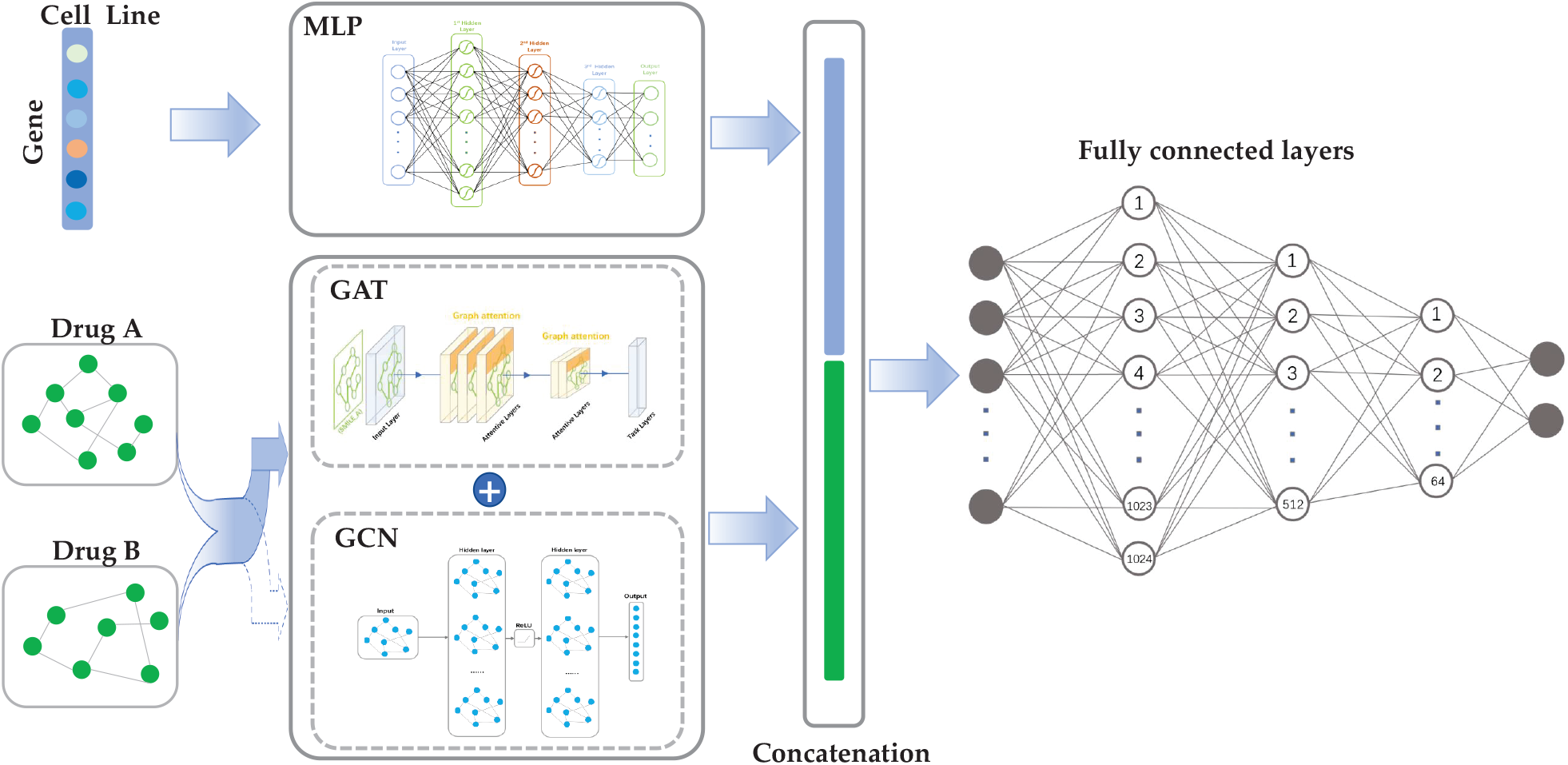
The pipeline of DeepDDS learning framework. The feature embedding of gene expression profiles of cancer cell line is obtained through Multi-Layer Perception (MLP), and the feature embedding of drug is obtained through GAT or GCN based on the drug molecular graph generated from drug SMILES. The embedding vectors of drug and cell line are subsequently concatenated to feed into a multi-layer fully connected network to predict the synergistic effect.

### Drug Representation based on GNN

We use the open-source chemical informatics software RDKit (Landrum et al., 2006) to convert the SMILES into molecular graphs, where the nodes are atoms and the edges are chemical bonds. Specifically, a graph for a given drug is defined as *G* = (*V, E*), where *V* is the set of *N* nodes that represented by a *C*-dimensional vector, and *E* is the set of edges represented as an adjacency matrix *A*. In a molecule graph, *x*_*i*_ ∈ *V* is the *i*-th atom and *e*_*ij*_ ∈ *E* is the chemical bond between the *i*-th and *j*-th atoms. The chemical molecular graph is non-Euclidean data and lacks of translation invariance, therefore, we applied graph neural network instead of traditional convolution network, to extract drug feature representations based on the graphs.

For each node in a graph, we use DeepChem (Ramsundar et al., 2019) to compute a set of atomic attributes as its initial feature. Specifically, each node is represented as a binary vector including five pieces of information: the atom symbol, the number of adjacent atoms, the number of adjacent hydrogen, the implicit value of the atom, and whether the atom is in an aromatic structure. In GNN, the learning process of drug representation is actually the message passing between each node and its neighbor nodes. In this paper, we consider two types of GNN (graph convolution network and graph attention network) in our learning framework and evaluate their performance in the drug feature extraction.

### Graph Convolutional Network (GCN)

The input of the multi-layer GCN is the node feature matrix *X* ∈ ℝ^*N* ×*C*^ and the adjacency matrix *A* ∈ ℝ^*N* ×*N*^ that represents the connection of nodes. According to Welling et al. (Kipf and Welling, 2016), it can write dissemination rules in a standardized format to ensure stability.

The iteration process can be defined as below:

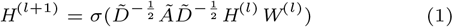

where *Ã* = *Ã* + *I*_*N*_ (*I*_*N*_ is the identity matrix) is the adjacency matrix of the undirected graph with added self-connections, 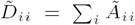 ; *H*^(*l*+1)^ ∈ ℝ^*N* ×*C*^ is the matrix of activation in the *l*th layer, *H*^(0)^ = *X, σ* is an activation function, and *W* is a learnable parameter.

The output *Z* ∈ ℝ^*N* ×*F*^ (*F* is the number of output features per node) can be defined as below:

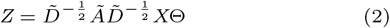

where Θ ∈ ℝ^*C*×*F*^ (*F* is the number of filters or feature maps) is the matrix of filter parameters.

Our GCN-based model uses three consecutive GCN layers activated by ReLU function. The original GCN is a method for classify the node by semi-supervised learning, i.e., the outputs are the node-level feature vectors. To construct graph-level feature vectors, we use Sum, Average, and Max Pooling to aggregate the whole graph feature from learned node features and evaluate their performance. We find that the use of Max Pooling layer in GCN-based DeepDDS outperforms the others. Therefore, we add a global Max Pooling layer after the last GCN layer to extract the representation.

### Graph Attention Network (GAT)

The graph attention network (GAT) proposes a multi-head attention-based architecture to learn higher-level features of nodes in a graph by applying a self-attention mechanism. Every attention head has its own parameters. The GAT architecture is built from the graphics attention layer. The output features for nodes were computed as

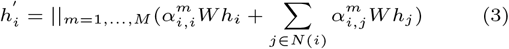

Where || concat the output results of multiple attention mechanisms, *M* is the number of attetion heads, and *W* ∈ ℝ^*C*′ ×*C*^ is a weight matrix. The attention coefficient *α*_*i,j*_, between each input node *i* and its first-order neighbor in the graph, is calculated as follows:

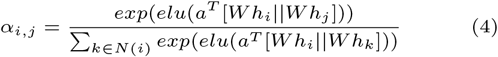

where *a*^*T*^ ∈ ℝ^*C*′^ is learnable weight vector, *T* is the corresponding transpose, and *elu* is a Non-linear activation function, when *x* is negative, *y* is equal to 0. Then, ‘softmax’ function is introduced to normalize all neighbor nodes *j* of *i* for easy calculation and comparison.

### Cell Line Feature Extraction based on MLP

To alleviate the dimension imbalance between the feature vectors of drugs and cell lines, we selected the significant genes according to a LINCS project (Yang et al., 2012). The LINCS project provides a set of about 1000 carefully chosen genes, referred to as ‘Landmark gene set’, which can capture 80% of the information based on the Connectivity Map (CMap) data (Cheng and Li, 2016). The intersected genes between the CCLE gene expression profiles and the Landmark set was chosen for subsequent analysis. We used the gene annotation information in the Cancer Cell Line Encyclopedia (CCLE) (Barretina et al., 2012) and the GENCODE annotation database (Derrien et al., 2012) to remove the redundant data, as well as the transcripts of non-coding RNA. Finally, we select **954** genes from raw expression profiles as input to the model.

We adopt an MLP to extract the cell line features. The MLP includes two hidden layers, and the number of hidden units of each layer is selected via hyperparameter selection (See Hyperparameter setting for detail).

### Predicting the synergistic effect of drug combinations versus cell lines

We formulated the prediction of synergistic drug combinations as an end-to-end binary classification model. Upon the embedding vectors of drugs through GAT or GCN, and the embedding vectors of cell lines through MLP, they are concatenated as the input of multiple fully-connected layers. We adopt the spindle-shaped structure for the fully connected layer. The probability of the synergistic effect (classification label) was computed by the softmax function that follows the output of the last hidden layer, as follows:

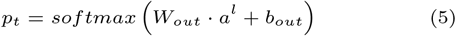

where *p*_*t*_ is the probability of t, *W*_*out*_ and *b*_*out*_ are the weight matrix and bias vector, *a*^*l*^ are the embedding features learned by previous layers, as follows:

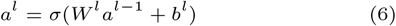

Where *l* is the number of hidden layers, *W* and *b* are the matrices corresponding to all hidden layers and output layers, bias vector, *a*^0^ = *concat*(*R*_*drug*1_, *R*_*drug*2_, *R*_*cellline*_) is the raw input vector.

Given a set of combinations with labels, we adopted the cross-entropy as the loss function to train the model, with the aim to minimize the loss during the training process, which is formulated as follows:

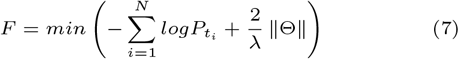

where Θ represents the set of all trainable weight and bias parameters involved in the model, *N* is the total number of samples in the training dataset, *t*_*i*_ is the *i*th label, and *λ* is an L2 regularization hyper-parameter.

## Result

### Hyperparameter setting

The real architecture of DeepDDS is actually determined by hyperparameter setting. The hyperparameters cover the numbers of layers and units of each layer in MLP, GCN and GAN, as well as the activation function and learning rate. As exhaustive enumerations of the hyperparameters are computationally inhibitive, thereby we adopt grid-like search to tune the hyperparameters. As shown in Table 1, we have tested different structural forms and values of these hyperparameters. We tuned the hyperparameters via five-fold cross validations on benchmark dataset. The selected values of these hyperparameters are displayed in boldface. The GCN yield to better performance in the drug feature extraction when its structure has three hidden layers and number of units are 1024, 512 and 156, respectively. We have also considered different number of hidden layers for GAN and MLP, and found they performed best with two hidden layers. For multihead attention mechanism, multiple independent values are evaluated. For the activation function, the ELU and ReLU activation functions after the GAT layers at DeepDDS-GAT are used. For DeepDDS-GCN, it also has similar layer structure, but only ReLU is used as activation function.

**Table 1.**
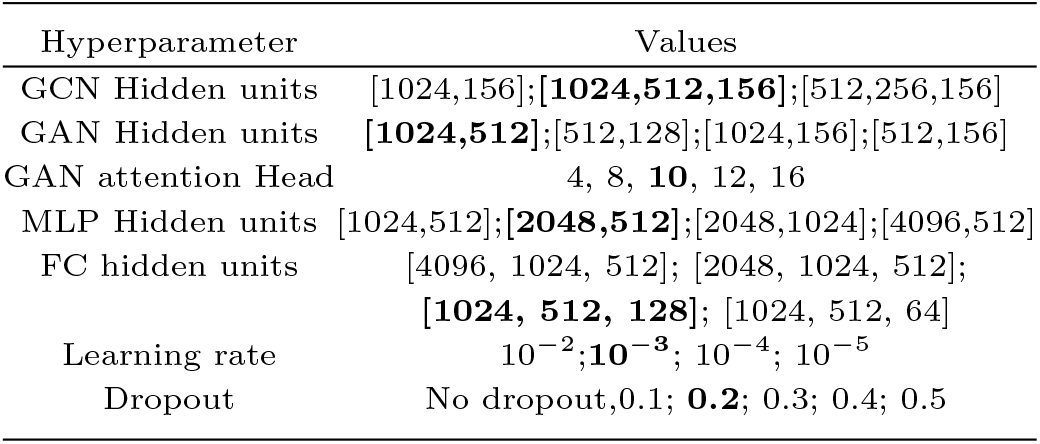
Hyperparameter settings of DeepDDS

| Hyperparameter | Values |
| --- | --- |
| GCN Hidden units | [1024,156];[**1024,512,156**];[512,256,156] |
| GAN Hidden units | [**1024,512**];[512,128];[1024,156];[512,156] |
| GAN attention Head | 4, 8, **10**, 12, 16 |
| MLP Hidden units | [1024,512];[**2048,512**];[2048,1024];[4096,512] |
| FC hidden units | [4096, 1024, 512]; [2048, 1024, 512];<br>[**1024, 512, 128**]; [1024, 512, 64] |
| Learning rate | $10^{-2}$; **$10^{-3}$**; $10^{-4}$; $10^{-5}$ |
| Dropout | No dropout, 0.1; **0.2**; 0.3; 0.4; 0.5 |

### Performance comparison on cross-validation

To evaluate the performance of DeepDDS, we compared DeepDDS with some current state-of-the-art methods, including both classical machine learning methods and deep learning-based methods. Six classical machine learning methods, including Random Forests (RF), Gradient Boosting Machines (GBM), Extreme Gradient Boosting (XGBoost), Adaboost, Multilayer Perceptron (MLP), Support Vector Machines (SVM), are considered in the performance comparison. Three deep learning-based methods are TranSynergy (Liu and Xie, 2021), DeepSynergy (Preuer et al., 2018) and Deep Tensor Factorization (Sun et al., 2020). To clarify the difference between DeepDDS and these deep learning-based methods, we summarize them as below:

- **TranSynergy**. TranSynergy includes three major components, input dimension reduction component, self-attention transformer component, and output fully connected component. It combines the network propagated drug target profile, gene dependency and gene expression to find novel genes associated with the synergistic drug combination from the learned biological relations.
- **DeepSynergy**. DeepSynergy uses molecular chemistry and cell line genomic information as input, and a cone layer in a neural network (DNN) to simulate drug synergy and finally predict the synergy score.
- **Deep Tensor Factorization (DTF)**. DTF combine tensor-based framework and deep learning methods together to predict synergistic effect of drug pairs, which is comprised mainly by a tensor factorization method and a deep neural network.

First, we conducted five-fold cross validation to benchmark the predictive power of DeepDDS. The training samples (each sample is a drug-drug-cell line triplet) are randomly split into five subsets of roughly equal size, each subset is taken in turn as a test set and the remaining four subsets are used to train the model, whose prediction accuracy on the test set is then evaluated. The average prediction accuracy over the 5-folds is used as the final performance measure. For clarity, we provide typical performance measures widely used in classification tasks, including area under the receiver operator characteristics curve (ROC AUC), area under the precision recall curve (PR AUC), accuracy (ACC), balanced accuracy (BACC), precision (PREC), sensitivity (TPR) and Cohen Kappa. Table 2 shows these performance measures of DeepDDS and other methods. Clearly, DeepDDS-GAT achieved higher accuracy than all other methods, and its performance measures of ROC AUC, PR AUC, ACC, BACC, PREC, TPR, TNR and Kappa reach 0.93, 0.93, 0.85, 0.85, 0.85, 0.85, 0.85 and 0.71, respectively. In fact, both DeepDDS-GAT and DeepDDS-GCN outperform others in terms of all these performance measures. We note that the classifier XGBoost also achieved remarkable performance, nevertheless still inferior to DeepDDS. The three deep learning-based methods TranSynergy, DTF and DeepSynergy follow closely XGBoost, but outperform other methods.

**Table 2.** Performance comparison of DeepDDS and competitive methods on 5-fold cross-validation

| Performance Metric | ROC AUC | PR AUC | ACC | BACC | PREC | TPR | KAPPA |
| --- | --- | --- | --- | --- | --- | --- | --- |
| DeepDDS-GAT | **0.93 ± 0.01** | **0.93 ± 0.01** | **0.85 ± 0.07** | **0.85 ± 0.07** | **0.85 ± 0.07** | **0.85 ± 0.07** | **0.71 ± 0.21** |
| DeepDDS-GCN | **0.93 ± 0.01** | **0.92 ± 0.01** | **0.85 ± 0.01** | **0.85 ± 0.01** | **0.85 ± 0.01** | **0.84 ± 0.01** | **0.70 ± 0.22** |
| XGBoost | 0.92 ± 0.01 | 0.92 ± 0.01 | 0.83 ± 0.01 | 0.83 ± 0.01 | 0.84 ± 0.01 | 0.84 ± 0.01 | 0.68 ± 0.01 |
| Random Forest | 0.86 ± 0.02 | 0.85 ± 0.02 | 0.77 ± 0.01 | 0.77 ± 0.01 | 0.78 ± 0.02 | 0.74 ± 0.01 | 0.55 ± 0.04 |
| GBM | 0.85 ± 0.02 | 0.85 ± 0.01 | 0.76 ± 0.02 | 0.76 ± 0.02 | 0.77 ± 0.01 | 0.74 ± 0.01 | 0.53 ± 0.04 |
| Adaboost | 0.83 ± 0.01 | 0.83 ± 0.03 | 0.74 ± 0.01 | 0.74 ± 0.02 | 0.74 ± 0.02 | 0.72 ± 0.01 | 0.48 ± 0.03 |
| MLP | 0.65 ± 0.02 | 0.63 ± 0.05 | 0.56 ± 0.06 | 0.56 ± 0.05 | 0.54 ± 0.04 | 0.53 ± 0.22 | 0.12 ± 0.04 |
| SVM | 0.58 ± 0.01 | 0.56 ± 0.02 | 0.54 ± 0.01 | 0.54 ± 0.01 | 0.54 ± 0.01 | 0.51 ± 0.12 | 0.08 ± 0.04 |
| TranSynergy | 0.90 ± 0.01 | 0.89 ± 0.01 | 0.83 ± 0.01 | 0.83 ± 0.01 | 0.84 ± 0.01 | 0.80 ± 0.01 | 0.64 ± 0.01 |
| DTF | 0.89 ± 0.01 | 0.88 ± 0.01 | 0.81 ± 0.01 | 0.81 ± 0.01 | 0.82 ± 0.01 | 0.77 ± 0.03 | 0.63 ± 0.04 |
| DeepSynergy | 0.88 ± 0.01 | 0.87 ± 0.01 | 0.80 ± 0.01 | 0.80 ± 0.01 | 0.81 ± 0.01 | 0.75 ± 0.01 | 0.59 ± 0.05 |

We further checked the top 100 drug pairs with highest predicted synergy scores by DeepDDS-GAT (For detail see Supplementary Table S1), and found that 98 drug pairs have been experimentally validated to be synergistic combinations over different cancer cell lines.

### Performance evaluation by input permutation

We found that the higher the real synergy score, the higher the predictive score. After normalization of real synergy scores to [0, 1] region, we draw a scatter plot of the drug combinations with respect to the predicted and real synergy scores. As shown in Figure 2 (a), most points locate closely to the identity line. The Pearson correlation between the predicted synergy scores and real synergy scores reach 0.801. The results indicate our method achieve superior predictive accuracy.

**Fig. 2.**
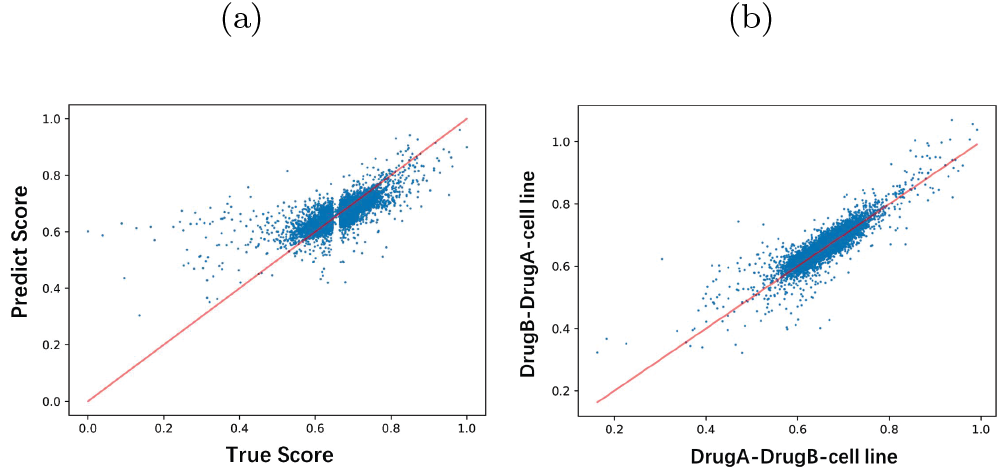
Scatter plots of synergy scores. (a) The scatter plot with respect to the real synergy scores and predicted synergy score. (b) The scatter plot of synergy score obtained from different input order of two drugs.

We go further to verify the predictive performance of DeepDDS upon different input order of two drugs. For drug A and drug B, we permutate the input features so that drug A-drug B and drug B-drug A are regarded as two different samples to train the model. Figure 2 (b) shows the predicted synergy score upon different sequence of input features by DeepDDS-GAT. It can be found that most values locate closely to the identity line and the Pearson correlation coefficient reach 0.9. It can prove that our model is insensitive to the sequence of the input features of drug combinations. In addition, we found that the ROC AUC and PR AUC obtained by drug A-drug B and drug B-drug A both reach or be close to 0.93.

### Performance evaluation by leave-one-out cross validation

We went further to verify the performance of the DeepDDS model using leave-one-out cross validation. First, we conducted the leave-one drug combination-out experiment. More precisely, we iteratively exclude each drug combination from the training set, use the remaining data to train the DeepDDS model that is in turn used to predict the sensitivity of the excluded drug combination to cancer cell lines. The result of the leave-one drug combination-out experiment is shown in Table 3, DeepDDS-GAT achieve notably performance by AUC value 0.89, followed by DeepDDS-GCN. It can be also found that DeepDDS significantly outperform all other methods.

**Table 3.**
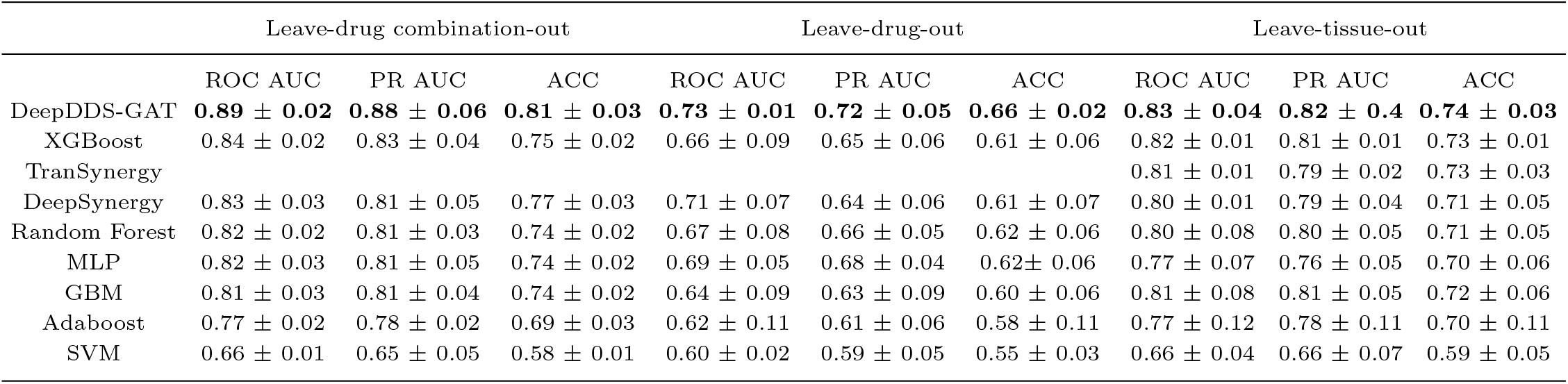
Performance on DeepDDS and competitive methods on leave-drug combination-out, leave-drug-out and leave-tissue-out experiments

As the leave-one drug combination-out experiment did not excluded single drug from the training set, we next leave one drug out to prevent the information of certain drug being seen by the model. The leave-one drug-out experiment check the potential to learn the important features of unseen drug from the chemical structures of those seen drugs. As shown in Table 3, DeepDDS still achieve better performance than other competitive methods.

As previous studies (Liu and Xie, 2021), we also carried out leave-one cell line-out experiment to verify the performance of DeepDDS. Take the cell line T47D as an example, the drug combination between BEZ-235 and MK-8669, Dasatinib, Lapatinib, Geldanamycin, PD325901, Erlotinib, MK-4541, Temozolomide, Vinorelbine, ABT-888, all have a high experimental synergy scores (Loewe>100). Expectedly, the prediction scores of these drug combinations have prior rankings among all candidate drug pairs (See Supplementary Table S2-S3 for detail). In addition to the leave-one cell line-out evaluation, Preuer et al., 2018), we adopt more rigorous strategy to evaluate our method. We exclude all the cancer cell lines belong to specific tissue from the training set, so that the model can not see any gene expression information of a certain type of tissue. We iteratively use the excluded cancer cell lines as the validation set and the remaining samples as the training set to train the model. Table 3 illustrated that DeepDDS-GAT achieve the best performance on leave-one tissue-out evaluation. Also, DeepDDS performs better than all classical machine learning methods and deep learning-based methods. Moreover, Figure 3 show the ROC AUC values of DeepDDS-GAT, DeepSynergy and TranSynergy on six different tissues, including breast, colon, lung, melanoma, ovarian and prostate. It can be found that DeepDDS-GAT is better than other two deep learning-based methods with ROC AUC 0.84, 0.867, 0.821, 0.828, 0.843 and 0.775 by leave-one tissue-out cross validation, respectively.

**Fig. 3.**
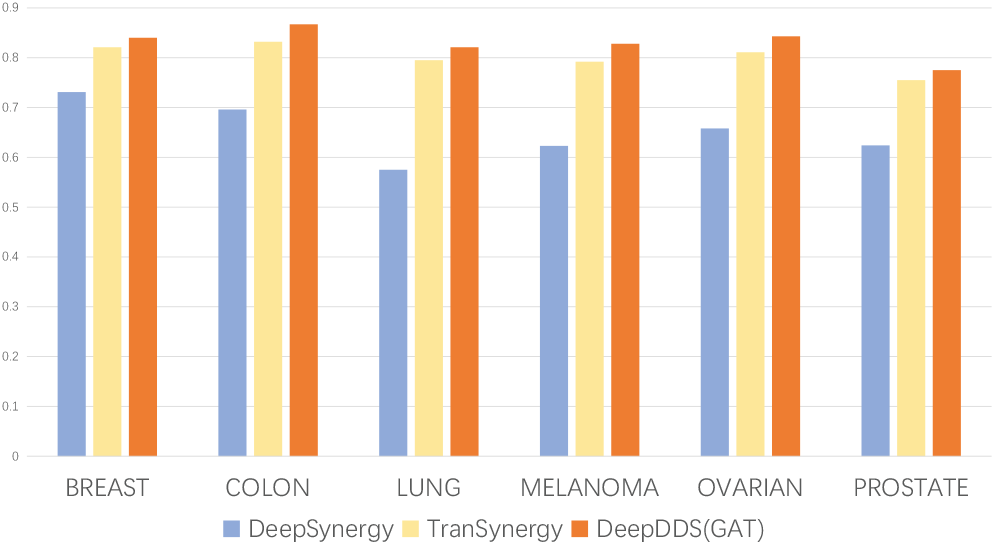
The ROC AUC values of DeepDDS-GAT, DeepSynergy and TranSynergy upon leave-tissue-out evaluations on six different tissues, including breast, colon, lung, melanoma, ovarian and prostate.

### Evaluation on independent test set

To verify the generalization ability of our method, we use the benchmark dataset (O’Neil et al., 2016) to train our model, and then employ an independent test set released by AstraZeneca (Menden et al., 2019) to evaluate the performance of DeepDDS and other competitive methods. The independent test set contains 668 unique drug-pair-cell-line combinations, covering 57 drugs (Supplementary Table S4) and 24 cell lines (Supplementary Table S5).

Table 4 shows the performance achieved by DeepDDS and competitive methods on the independent test set. It can be seen that the performance of DeepDDS is better than all competitive methods in terms of every performance measure. For clarity, we draw the ROC curves of DeepDDS and other methods, as shown in Figure 4. DeepDDS-GAT and DeepDDS-GCN account for top 2, followed by DeepSynergy. Meanwhile, it can be found that most machine learning-based methods perform just as random guess. This result indicate classical machine learning methods run into overfitting, while deep learning-based method acquire better generalization ability. In particular, DeepDDS-GAT and DeepDDS-GCN correctly predicted 421 (421/668=0.63) and 402 (402/668=0.6) drug pairs included in the independent test set, which outperform DeepSynergy correct prediction 317 (317/668=0.47) by 16% and 13%, respectively. The confusion matrices in Figure S3 show detailed numbers of correctly and falsedly predicted samples by the three methods.

**Table 4.**
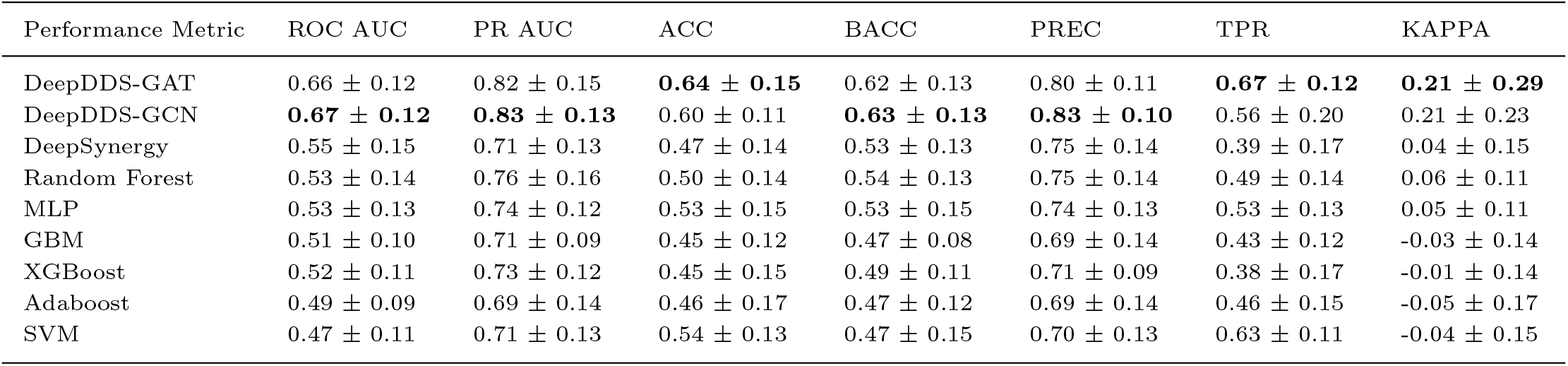
Performance metrics for the classification task in independent test set

**Fig. 4.**
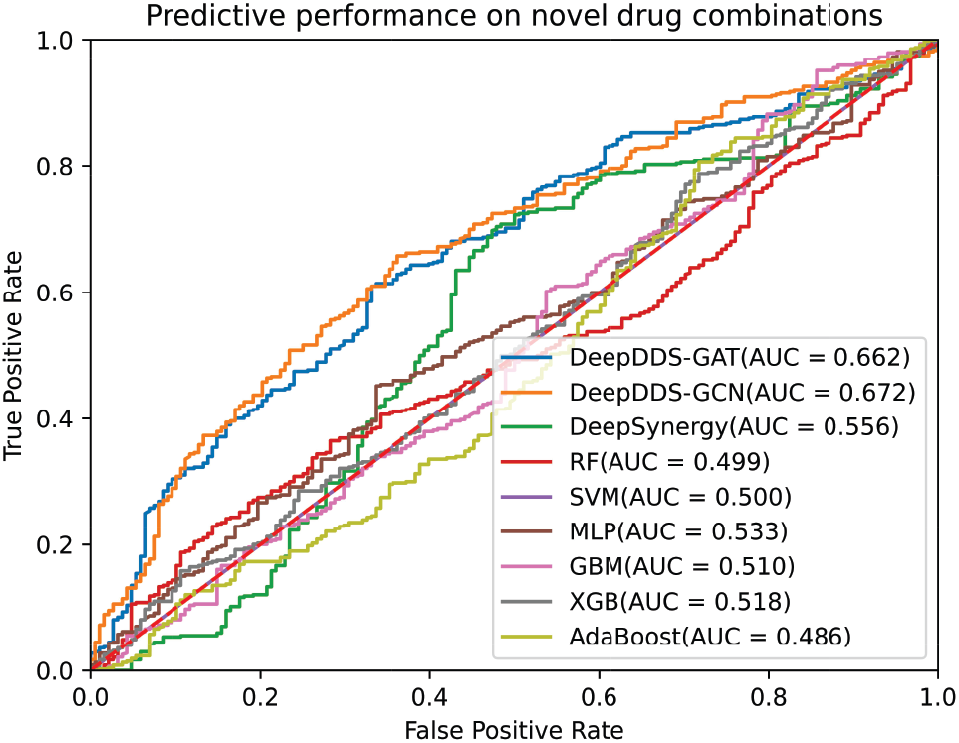
ROC curves and AUC values of DeepDDS and competitive methods on independent test dataset released by AstraZeneca.

### Graph attention network reveals important chemical substructure

DeepDDS-GAT model iteratively passes messages between nodes so that each node can capture the information of its neighboring nodes. Meanwhile, each neuron is connected to the neighborhood upper layer through a set of learnable weights in the GAT network. As a result, the feature representation actually encodes the information of the chemical substructure around the atom, including formal charge, water solubility and other physicochemical properties. This motivate us to explore the implications of the attention mechanism in revealing the important chemical substructures.

For example, previous study has showed that EGFR inhibitor Afatinib and AKT inhibitor MK2206 play synergistic effect in the treatment of lung cancer, head and neck squamous cell carcinoma (HNSCC) (Modjtahedi et al., 2014,Silva-Oliveira et al., 2017,Hung et al., 2016). We investigate how the atomic feature vectors evolved during the learning process, by measuring the Pearson correlation coefficient between atom pairs based on the feature vectors. The heat maps of the atom correlation matrix is plotted to observe the change of feature patterns. The similarity scores are displayed in the cells and indicated by the color scheme. It can be seen that before training the visual patterns in the heat maps of two drugs shows some degree of chaos. After training, however, the heat map of both drugs show obvious atomic clusters in a specific order. In particular, the atom of drug Afatinib is clustered into five subgroups, while MK2206 clustered into two atom subgroups (one big and one small block), as shown in figure 5. Without loss of generalization, we randomly select a few other drug combinations to check whether their feature vectors undergo similar pattern changes during the training process. These drug combinations include AZD2014 and AZD6244, AZD8931 and AZD5363, GDC0941 and AZD6244,GDC0941 and MK2206. As expected, the atomic feature vectors of the involved drugs gradually cluster into several subgroups (See Supplementary Figure S2-S6 for more detail).

**Fig. 5.**
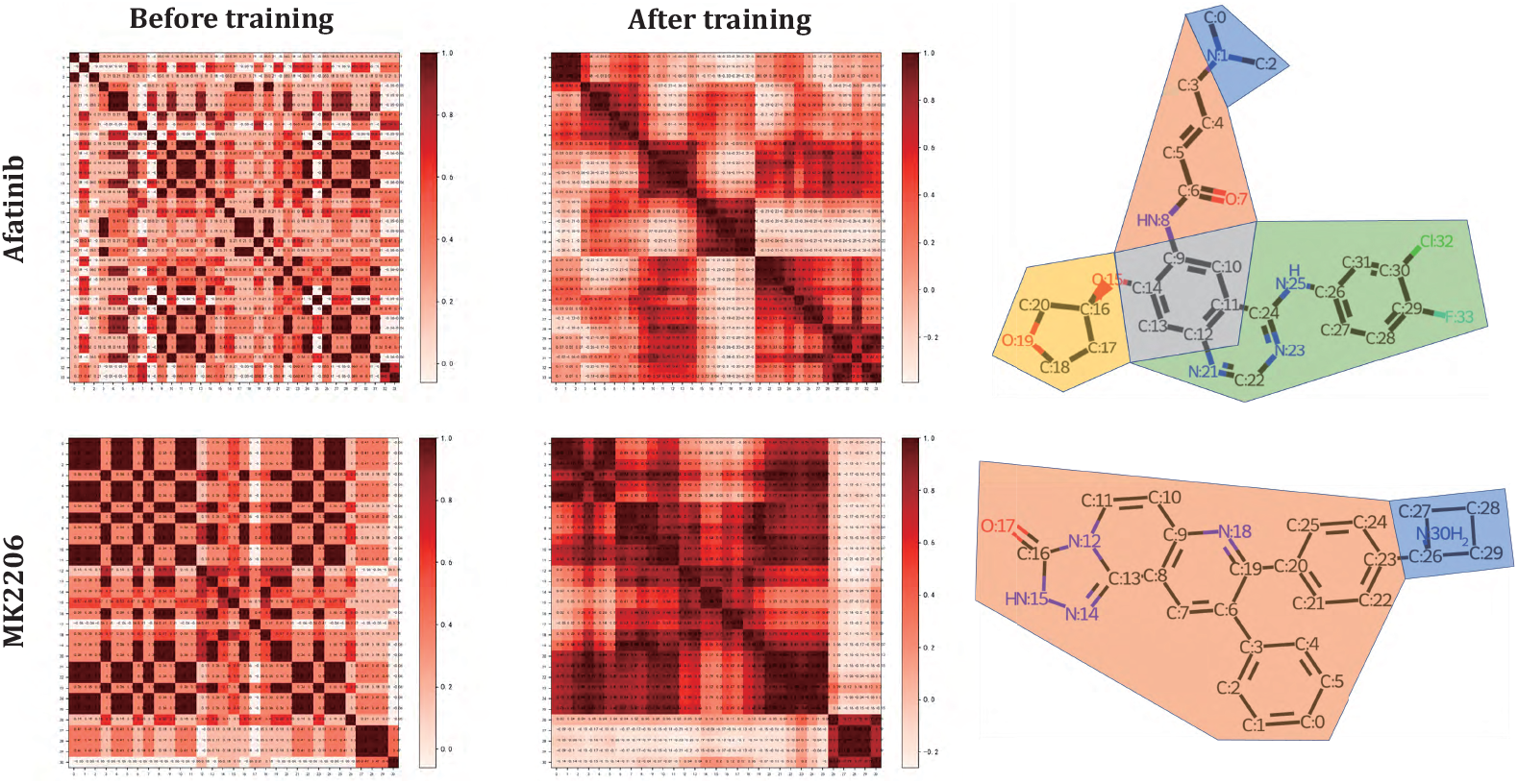
Heat map of the atomic feature similarity matrices of Afatinib and MK2206 before and after training. The heat maps show clear clustering patterns during the learning process. The diagrams of chemical structures of Afatinib and MK2206 display the five and two subgroups according to their clusters of heat maps.

We go a step further to explore the interpretability of the graph attention network in revealing the chemical substructures that are potential components exerting synergistic effect of the drug combinations. We compute the Pearson correlation coefficients between atom pairs across two drugs, so that significant association between chemical subgroups of different drugs can be uncovered. Take the drug combinations Afatinib and MK2206 as example again, we find that the heat map of the atom correlation matrix have no clear clustering pattern before training, while it show two notably linking blocks after training, as shown in Figure 6 (a). More interesting, these two linking blocks exactly indicate that the bigger atom subgroup (No.1-25 atoms) of MK2206 associates to the 3th and 5th atom subgroups (No.9-14 atoms and No.21-33 atoms) of Afatinib. From the 3D structures of the two drugs, they are just the main functional groups of Afatinib and MK2206, respectively. For other examples mentioned above, we found that their inter-drug atom correlation matrices also display clustering patterns, as shown in Figure S3-S6.

**Fig. 6.**
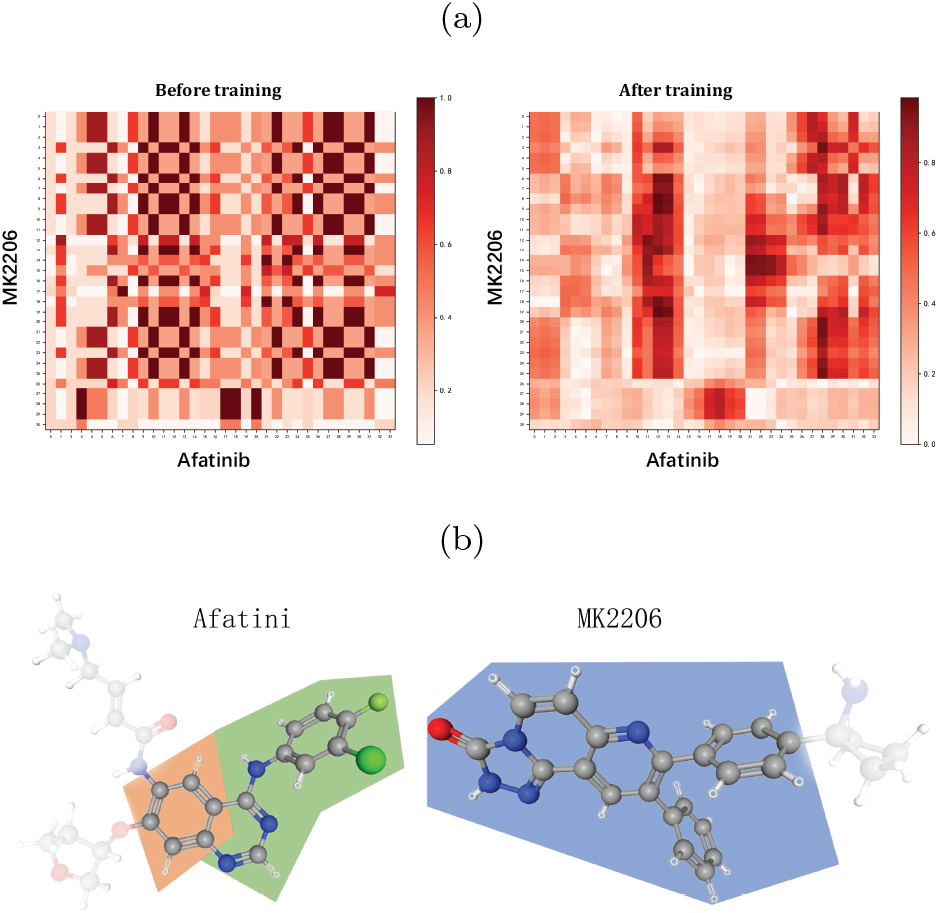
The heat maps of Pearson correlation coefficients between atom pairs across Afatinib and MK2206. Pearson correlation coefficients are computed using the feature vector before and after training. (a) The heat map shows no clear visual pattern before training, but after training shows two clear linking blocks. (b) The bigger atom subgroup (No.1-25 atoms) of MK2206 associates remarkably to the 3th and 5th atom subgroups (No.9-14 atoms and No.21-33 atoms) of Afatinib.

As a result, the atom embedding vectors display clear feature patterns during the training process, namely, the atom correlation matrices clearly cluster into several atom subgroups, and the degree of association between atom subgroups of different drugs transfer from chaos to order. We adventure to speculate that the atom subgroups included in these two drugs play key role in their synergistic function, although the pharmacological mechanism in vivo remains unclear to date.

### Predicting novel synergistic combinations

The performance evaluation experiments above have shown that our DeepDDS model achieve superior performance, thereby we apply DeepDDS to predict novel synergistic combination. We use the O’Neil drug combination dataset to train the DeepDDS model. To generate candidate drug combinations, we selected 42 small molecule targeted drugs approved by the FDA (Bedard et al., 2020) and then generated 855 candidate drug pairs (see Supplementary Table S6). We listed the top 10 predicted drug combinations in Table 5. To verify the reliability of the predicted results, we conducted an non-exhaustive literature search and found there are at least six predicted drug combinations are consistent with the observations in previous studies or under clinical trials. We presented the PMIDs or DOI identifiers of these related publications in Table 5.

**Table 5.** Top 10 predicted novel synergistic combinations on A375 cancer cell line

| Drug A | Drug B | Cell line | Predict Score | Publications |
| --- | --- | --- | --- | --- |
| **Abemaciclib** | **Lapatinib** | A375 | 0.9977 | 26977878, 33389550, 26977873 |
| Binimetinib | Sorafenib | A375 | 0.9974 | NA |
| Copanlisib | Regorafenib | A375 | 0.9973 | NA |
| **Copanlisib** | **Sorafenib** | A375 | 0.9973 | 30962952, 27259258, doi:10.5282/edoc.24304 |
| Binimetinib | Regorafenib | A375 | 0.9971 | NA |
| **Erlotinib** | **Regorafenib** | A375 | 0.997 | 25907508 |
| **Vemurafenib** | **Sorafenib** | A375 | 0.9969 | 33119140, 30076136, 30844744, 29605720, doi:10.21037/tcr.2020.01.62 |
| Vemurafenib | Regorafenib | A375 | 0.9967 | NA |
| **Lapatinib** | **Regorafenib** | A375 | 0.9967 | 27864115, 24911215 |
| Pazopanib | Sorafenib | A375 | 0.9965 | NA |

For example, the CDK4/6 inhibitor **abemaciclib** and EGFR inhibitor **lapatinib** significantly enhanced growth-inhibitory for HER2-positive breast cancer (Goel et al., 2016). Ye *et al*. found that **Copanlisib** reduced **Sorafenib**-induced phosphorylation of p-AKT and enhanced synergistically of antineoplastic effect in vitro (Ye et al., 2019). Also, the combination of **Erlotinib** and **Regorafenib** in the treatment of hepatocellular carcinoma successfully overcome the interference of epidermal growth factors (DAlessandro et al., 2015). Addition of **Sorafenib** to **Vemurafenib** increased ROS production through ferroptosis, thus increasing the sensitivity of melanoma cells to vemurafenib (Tang et al., 2020). Zhang et al. reproted that the **Regorafenib** combined with the **Lapatinib** could improve anti-tumor efficacy in human colorectal cancer (Zhang et al., 2017). We believe that other predicted drug pairs are also promising combinations await for further validation.

## Discussion and Conclusion

In this paper, we have proposed a novel method to predict synergistic drug combination to specific cancer cells. Overall, our method performs significantly better than other competitive methods on the five-fold cross validation experiments. However, we noticed that the predictive accuracy of our method are still limited on the independent test set, although the performance of our method is greatly superior than all competitive methods. We think the limited performance is mainly attributed to the small number of training samples. In fact, the benchmark dataset actually includes only 38 unique drugs and 39 cancer cell lines, while the space for possible drug combinations is much larger when novel drugs are included

Two different graph neural network, GAT and GCN, are used to learn drug embedding vectors in our method. We extensively compared their performance to each other, as well as to quite a few competitive methods. Overall, GAT performs slightly better than GCN, and thus we further explored the interpretability of the GAT model. However, we have realized that the physicochemical properties of the molecular graph and attention weights between the atoms have not been fully understood. In the future, we are interested in studying the connections between atoms to incorporate more information resources into the DeepDDS model to improve the model interpretability and predictability.

In conclusion, we have proposed a novel method DeepDDS to predict the synergy of drug combinations for cancer cell lines with high accuracy. Our performance comparison experiments showed that DeepDDS performs better than other competitive methods. We have demonstrate that DeepDDS achieve state-of-the-art performance in a cross-validation setting with an independent test set. We believe that with the increasing size of the data set available, DeepDDS can be further improved and applied to other fields where drug combinations play an essential role, such as antiviral (Akhtar, 2020), antifungal (Pereira et al., 2021) and multi-drug synergy prediction(Ontong et al., 2021). Overall, we believe that our method will yield some inspiring insights into the discovery of synergistic drug combinations.

## Author contributions statement

J.W. and H.L. conceived the main idea and the framework of the manuscript. J.W. drafted the manuscript. J.W. and X. L. collected the data and performed the experiments. L.D. and H.L. helped to improve the idea and the manuscript. S.S. reviewed drafts of the paper. L.D. and H.L. supervised the study and provided funding. All authors read and commented on the manuscript.

## Acknowledgments

This work was supported by the National Natural Science Foundation of China under grants No. 61972422 and No. 62072058.

**Jinxian Wang**. Jinxian Wang received the Bachelor’s degree from hunan Agricultural University in 2019, and at present is studying for a masters degree at Central South University supervised by Prof. Lei Deng. His study focuses on machine learning and bioinformatics.

**Xuejun Liu**. Xuejun Liu is a professor at School of Computer Science and Technology, Nanjing Tech University, Nanjing, China. His research interests include data mining and deep learning.

**Siyuan Shen**. Siyuan Shen is a graduate student at School of Software, Xinjiang University, Urumqi, China. His research interest is using machine learning algorithms to study non-coding RNA interactions and functions.

**Lei Deng**. Lei Deng is a professor at School of Computer Science and Engineering, Central South University, Changsha, China. His research interests include data mining, bioinformatics and systems biology.

**Hui Liu**. Hui Liu is a professor at School of Computer Science and Technology, Nanjing Tech University, Nanjing, China. His research interests include the anti-cancer drug screening by means of bioinformatics and deep learning.

